# Protein without farms: What comparative genomics reveals about “Power-to-Food” microbes

**DOI:** 10.64898/2026.04.04.716520

**Authors:** Karan Kumar, Juha-Pekka Pitkänen, Lars M. Blank, Tobias B. Alter

## Abstract

Conventional agriculture is increasingly incompatible with planetary boundaries, such as land and water demand, greenhouse-gas emissions, and disruption of the nitrogen cycle. Hydrogen-oxidizing bacteria (HOB) enable a scalable “power-to-food” approach in which aerobic gas fermentation turns CO₂ and renewable H₂, along with N₂ in some strains, into protein-rich biomass, largely decoupling protein production from arable land and climate variability. The same chemistry is attractive for closed-loop space life support, where crew CO₂ and electrolysis-derived H₂ can be recycled into edible biomass. Here, we compare two leading HOB chassis strains, *Cupriavidus necator* H16 and *Xanthobacter* sp. SoF1, using standardized re-annotation, orthology-based comparison, pathway reconstruction, and safety-oriented genome screening. Importantly, SoF1 is the production strain for Solar Foods’ Solein®, a dried microbial biomass ingredient, which is approved as a novel food in Singapore and has a self-affirmed GRAS status in the United States. H16 has a larger, multipartite genome of 7.41 Mb split across two chromosomes and the pHG1 megaplasmid, whereas SoF1 is more compact at 4.91 Mb and encoded on a single replicon. Both encode Calvin–Benson–Bassham CO₂ fixation and multiple [NiFe]-hydrogenase systems supporting growth on CO₂/H₂, but nitrogen economy differentiates the hosts. SoF1 encodes a complete nitrogen-fixation module (*nifHDK*) and nitrate-assimilation genes, whereas H16 lacks *nif* and instead encodes nitrate/nitrite respiration for oxygen-limited flexibility. Safety screening revealed no evidence of canonical virulence determinants, integron or plasmid-linked antimicrobial resistant (AMR) cassettes, or high-confidence foodborne exotoxins under strict thresholds. These results convert genome-level features into actionable design constraints for selecting and engineering food-grade HOB, strengthening robust air-to-protein bioprocesses on Earth and informing a blueprint for closed-loop, space-compatible protein production.

**Highlights:**

- Hydrogen-oxidizing bacteria enable power-to-protein from CO₂, H₂ with minimal land use.
- Head-to-head genomics defines design rules for food-grade “air-to-protein” bioprocesses.
- Contrasting nitrogen routes guide media design, nutrient inputs, and closed-loop operation.
- CO₂ fixation and hydrogenase gene sets reveal complementary robustness and control features.
- Genome architecture and COG shifts inform safety, stability, and regulatory-ready strain choice.

## Background

Food production via traditional agricultural practices is planetary boundaries. The planetary health check 2025 report reveals that seven of the nine critical Earth-system boundaries are already transgressed, including climate change and biogeochemical flows.^1,2^ In particular, anthropogenic reactive nitrogen inputs exceed the proposed global boundary (∼62 Tg N yr⁻¹), reflecting the scale of fertilizer-driven disruption underpinning modern agriculture.^3^ At the same time, food systems account for one third of global anthropogenic greenhouse-gas emissions (2018 estimate), largely from agriculture and land-use change, intensifying pressure to produce protein with radically lower land, water, and fertilizer demands.^4,5^ These constraints collide with demographics: updated UN projections place the global population near ∼9.7 billion by 2050, increasing demand for affordable, high-quality protein without further expanding cropland or nitrogen pollution.^6^

To meet rising protein demand sustainably, alternative protein sources such as microbial single-cell protein are gaining again attention.^7^ Microbial biomass can be extremely protein-rich (30–80% dry weight) and cultivated without arable land.^8^ In particular, certain chemoautotrophic bacteria such as hydrogen-oxidizing bacteria (HOB) offer a compelling “power-to-food” alternative.^6,9^ In gas fermentation, aerobic HOB convert CO₂ and renewable H₂ (and, for some strains, N₂) into protein-rich biomass, enabling food production largely independent of arable land and climate variability.^6,10^ This premise of turning crew CO₂ and recycled resources into edible biomass also aligns with the logic of closed-loop life support of NASA and other space agencies.^11,12^ They have long emphasized bioregenerative systems that recycle wastes into oxygen, water, and food to reduce resupply mass and improve space mission autonomy.^11^ HOB-based processes are attractive in such closed loops because their core inputs (CO₂, H₂, minerals) can be generated from onboard recycling plus electrolysis powered by solar or nuclear energy.^7,13^

HOB such as *Cupriavidus necator* H16 (hereafter H16) and *Xanthobacter* sp. SoF1 (hereafter SoF1) grow on CO₂, H₂, O₂, and mineral nutrients, converting renewable energy into edible biomass.^8^ Their cells contain ∼60-70% protein with an amino acid profile comparable to animal protein.^9^ Yet, widespread adoption of microbial biomass as food remains gated by consumer acceptance and regulatory proof of safety, compositional consistency, and process control under distinct regional regimes.^7,8,14^ In the European Union, microbial biomass typically falls under the Novel Food framework (Regulation (EU) 2015/2283)^15^ requiring European Food Safety Authority safety evaluation^16^ and Commission authorization; in the US, the GRAS notification program provides a voluntary pathway to document safety conclusions and obtain FDA feedback.^16,17^

Within this context, H16 is a deeply characterized hydrogenotroph historically explored for single-cell protein production^8^ and is now re-emerging as a chassis for sustainable biomanufacturing from C1 compounds.^18,19^ It was investigated as a single-cell protein source in the 1970s and can accumulate up to ∼70% protein in dry biomass under optimal conditions.^9,13^ Its complete genome sequence and extensive physiological characterization make it a valuable benchmark for systems-level analysis.^18^ In contrast, SoF1, the production strain used for Solar Foods’ Solein ®,^20,21^ is a newer HOB isolated from the seashore of the Baltic Sea in Finland.^22^ SoF1 represents an industrially relevant “air-to-protein” chassis supported by emerging safety data, including repeated-dose oral toxicity assessment of SoF1-derived biomass, that directly informs food-approval dossiers.^23,24^ In addition, SoF1 can fix atmospheric nitrogen, enabling growth without added ammonia and offering a potential advantage in resource-limited or closed-loop settings.^22^ Together, these two strains exemplify the promise of HOB as nutritious, low-footprint protein platforms for terrestrial and space-linked food systems.

In this study, H16 and SoF1 are selected for comparative genomic analysis because they represent two of the most relevant HOB chassis at the interface of sustainable biomanufacturing and food biotechnology. H16 as a deeply characterized model organism with a long history in single-cell protein and C1 bioprocess research,^19^ and SoF1 as an emerging industrial “air-to-protein” production strain already linked to food-application development. Despite this shared relevance, no comparative genomic analysis^25^ of H16 and SoF1 has yet been reported in the context of food biotechnology. A genome-level comparison is needed to resolve differences in metabolic pathways, carbon and nitrogen utilization, stress-response capacity, and safety-relevant traits that may directly affect protein yield, cultivation robustness, and regulatory assessment. Such a comparison is important to (i) rationalize strain selection for sustainable protein production, (ii) anticipate genome-encoded features relevant to safety and regulatory review, including undesired metabolite potential, mobile elements, and other risk-associated loci, and (iii) guide future engineering toward robust, food-grade, and closed-loop-compatible bioprocesses. We therefore present a comparative genomic survey of H16 and SoF1 to identify the genetic features that distinguish their capabilities and to inform the design of next-generation “air-to-food” platforms.

## Results

### Genome overview and relatedness

Although *Cupriavidus necator* H16 belongs to the Betaproteobacteria-related Burkholderiales group and *Xanthobacter* sp. SoF1 to the Alphaproteobacteria-related Xanthobacterales group, both converge on a shared hydrogen-oxidizing, chemolithoautotrophic chassis logic. H16 is a complete multipartite genome (RefSeq: GCF_004798725.1) comprising two chromosomes (Chr1 4,049,965 bp; Chr2 2,912,457 bp) and the 452,139 bp megaplasmid pHG1, totaling 7,414,561 bp with 66.3% GC. In contrast, SoF1 (strain deposit VTT-E-193585; internal assembly) is organized as a single replicon of 4,913,175 bp with a slightly higher GC content (67.8%). Table **1** presents an overview and general features comparison of both hosts. Despite a ∼2.5 Mb difference in genome size, both genomes exhibit similarly high coding density (H16 88.4% vs SoF1 87.2%) and a comparable fraction of protein-coding loci (CDSs: H16 6,741; 98.4% of genes; SoF1 4,434; 98.1%), with an identical proportion of proteins receiving functional predictions (93.8% in both). H16 encodes 6,853 genes and 84 RNA features, including 65 tRNAs and 15 rRNAs, whereas SoF1 contains 4,520 genes and 64 RNA features (50 tRNAs; 6 rRNAs) and a single CRISPR-associated locus. In addition, the H16 megaplasmid contributes 454 genes and is GC-depleted relative to the chromosomes (62.3% GC), consistent with a specialized accessory replicon. At the functional-category level, both genomes contain a large “function unknown” fraction (COG S ∼25–26%), and in the plotted COG subset (Fig. **1**), SoF1 shows higher representation of amino-acid transport and metabolism (COG E 13.2% vs 10.1% in H16), while H16 is higher in transcription (COG K 13.3% vs 9.2%) and energy production and conversion (COG C 12.9% vs 10.4%).

**Figure 1:**
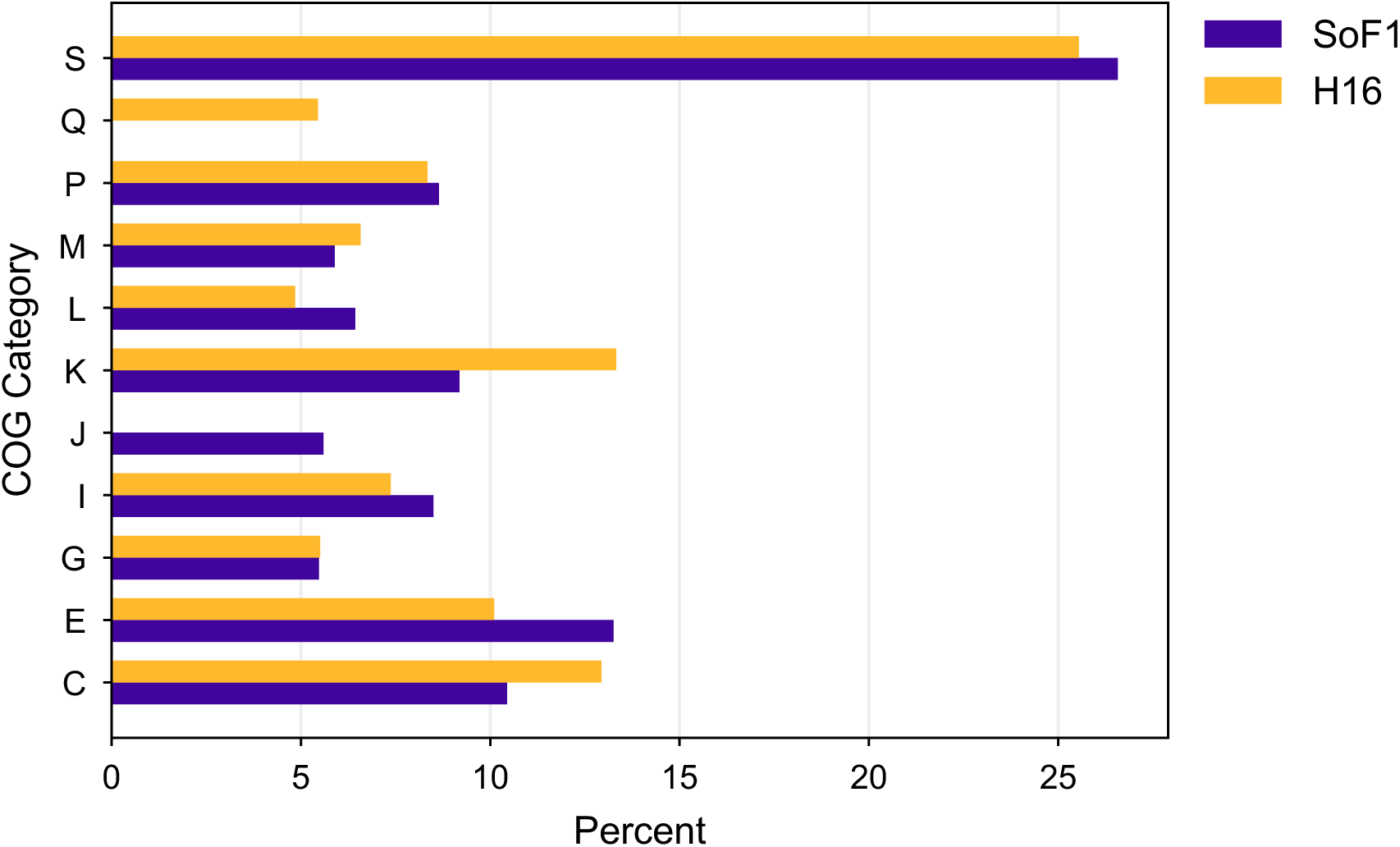
Functional composition of selected Clusters of Orthologous Groups (COG) categories in *Xanthobacter* sp. SoF1 and *Cupriavidus necator* H16. Bar chart comparing the relative abundance of selected COG functional categories among COG-assigned protein-coding genes in SoF1 and H16. Values represent the percentage of genes assigned to each plotted category: **C**, energy production and conversion; **E**, amino acid transport and metabolism; **G**, carbohydrate transport and metabolism; **I**, lipid transport and metabolism; **J**, translation, ribosomal structure and biogenesis; **K**, transcription; **L**, Replication, recombination and repair; **M**, cell wall/membrane/envelope biogenesis; **P**, inorganic ion transport and metabolism; **Q**, secondary metabolites biosynthesis, transport and catabolism; and **S**, function unknown. The profiles indicate overall similarity in core cellular functions, with notable differences in amino acid metabolism, transcription, and energy production or conversion.

**Table 1:** Overview of general features and metabolic pathway comparison of H16 and SoF1.

| Characteristics | *Cupriavidus necator* H16 |  | *Xanthobacter* sp. SoF1 |  |
| --- | --- | --- | --- | --- |
|  |  | Percent |  | Percent |
| Identifier / accession | GCF_004798725.1; Chr1<br>NZ_CP039287.1; Chr2<br>NZ_CP039288.1; pHG1<br>NZ_CP039289.1 |  | Strain deposit: VTT-E-193585; internal assembly (not publicly accessioned) |  |
| Replicons | 2 chromosomes + 1 plasmid (pHG1) |  | Single replicon |  |
| Chromosome 1 (bp) | 4,049,965 | 4 Mb |  |  |
| Chromosome 2 (bp) | 2,912,457 | 2.9 Mb |  |  |
| Megaplasmid pHG1 (bp) | 452,139 | 0.45 Mb |  |  |
| Total genome size (bp) | 7,414,561 (all replicons) | 7.4 Mb | 4,913,175 | 4.9 Mb |
| N50/N90 | N50 4,049,965; N90 2,912,457 (3 replicons) |  | 4,913,175 |  |
| Coding density | - | 88.4 | - | 87.2 |
| G+C composition % | - | 66.3 | - | 67.8 |
| Genes | 6821 |  | 4520 |  |
| Coding sequences (CDSs) | 6736 | 98.4 | 4434 | 98.1 |
| Protein genes with func. pred.* | 6430 | 93.8 | 4240 | 93.8 |
| CRISPR related genes | 0 | - | 1 | - |
| Total RNA | 85 | 1.4 | 64 | 1.5 |
| tRNA | 69 | 0.95 | 50 | 1.1 |
| tmRNA | 1 | - | 1 | - |
| rRNA | 15 | 0.21 | 6 | 0.13 |
| ncRNA | 17 | 0.24 | 11 | 0.24 |
| ncRNA regions (Regulatory) | 14 | 0.2 | 17 | 0.38 |
| Hypotheticals | 311 | 4.5 | 194 | 4.3 |
| Pseudogenes | 14 | - | 5 | - |
| Total no. of enzymes* | 1926 | 28.1 | 1913 | 36.4 |
| Enzymatic reactions* | 2624 | 28.1 | 2723 | 36.4 |
| Transporters* | 540 |  | 619 |  |
| Transport reactions* | 88 |  | 88 |  |
| Pathways | 488 |  | 527 |  |
| Putative HGT | 2 | - | 7 | - |
| COGs* | 6055 | 88.3 | 3955 | 87.5 |
| Plasmid No. of bases | 452,139 | 0.45 Mb | - | - |
| No. of genes | 429 | - | - | - |
| G+C % | - | 62.3 | - | - |
| Notes relevant to autotrophy | pHG1 carries major hydrogenase capacity and associated maturation/regulation modules in classic H16 literature |  | SoF1 autotrophy to CBB cycle and hydrogenase-driven energy metabolism |  |

\* including genes on plasmid

**Table 2:**
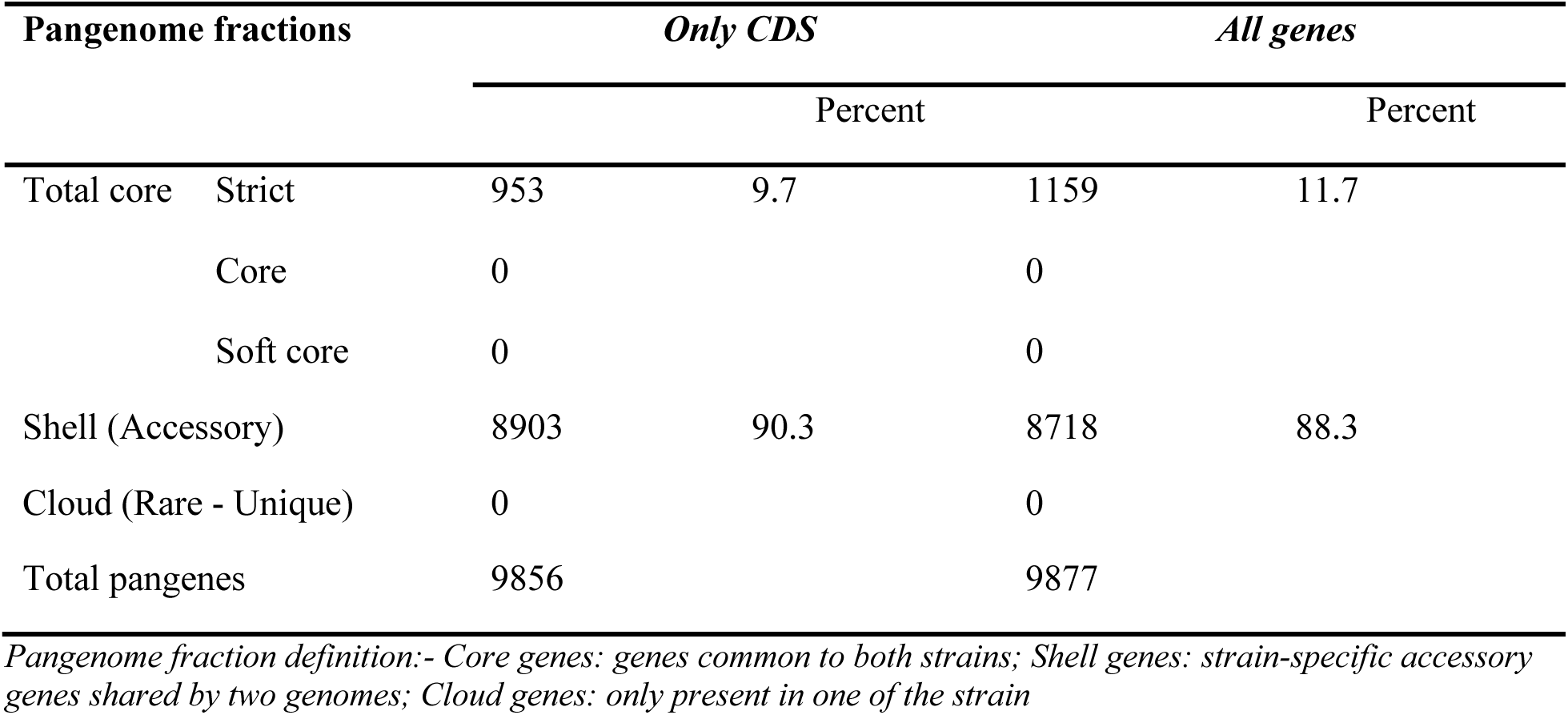
Pan-genome fractions in all genes and CDS-related.

Circos-based^26^ comparison resolves a conserved backbone between the single-replicon SoF1 chromosome and H16’s multipartite genome (Chr1, Chr2, and pHG1). Figure **2** summarizes the replicon-scale architecture and homology structure between SoF1 and the multipartite H16 genome. Although SoF1 carries a single chromosome and H16 is partitioned into two chromosomes plus pHG1, the genomes retain substantial shared protein-level content. The predominantly noncollinear correspondences, however, point to extensive rearrangements and translocations, including exchanges involving H16 chromosome 2 and pHG1.^27^ Feature overlays mark tRNA loci across each replicon and identify antiSMASH-predicted BGC intervals. Whereas the tRNA loci simply indicate the distribution of core translation-related elements, the BGC intervals occupy only a small fraction of the total genome span, suggesting that secondary-metabolite biosynthetic capacity is present in both strains but not extensive.^27^ Overall, the plot indicates that the two strains share a conserved core functional repertoire even though their genomes are organized differently. The limited genomic span occupied by antiSMASH-predicted BGCs in both strains further suggests that they are oriented toward efficient growth and redox/carbon metabolism rather than broad secondary-metabolite production.

**Figure 2:**
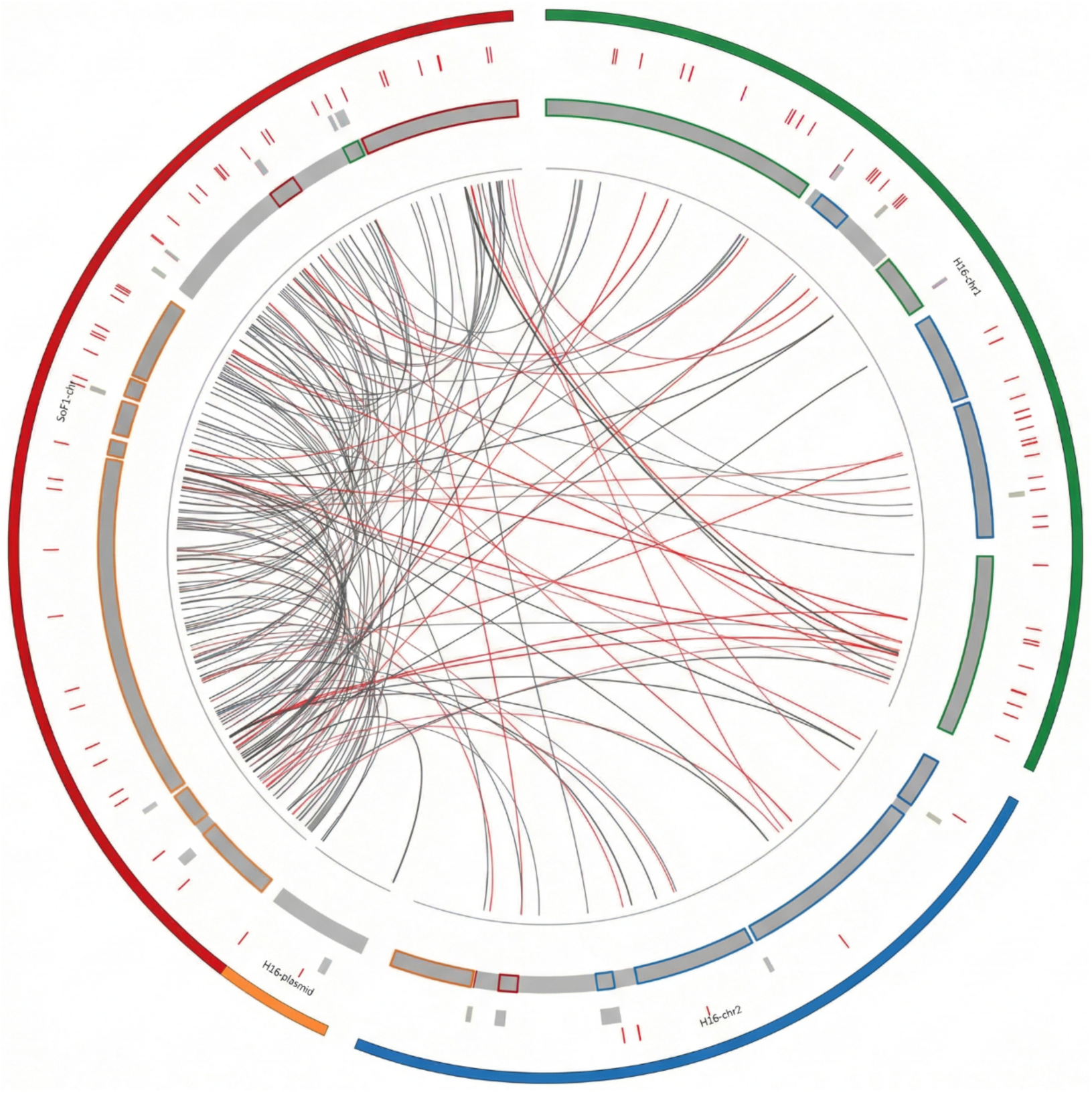
Genome architecture and homologous link map between H16 and SoF1. Circular ideograms (outermost colored arcs) represent replicons: SoF1 chromosome (red) and the multipartite H16 genome comprising chromosome 1 (green), chromosome 2 (blue), and the megaplasmid pHG1 (orange). The outer tick track marks tRNA loci mapped along each replicon. Predicted biosynthetic gene cluster (BGC) intervals from antiSMASH are shown as segmented blocks on the grey feature ring, with outline colors denoting BGC class. Interior ribbons connect homologous features across genomes: black links denote reciprocal best-hit ortholog anchors derived from DIAMOND protein alignments after quality filtering, while red links highlight a curated subset emphasizing non-collinear/synteny-disrupting correspondences. The dense backbone of links indicates extensive shared gene content, whereas the prevalence of crossing (non-collinear) ribbons is consistent with rearrangements and replicon-level re-partitioning superimposed on conserved segments.

### Core-accessory genome partitioning

PEPPAN-based pangenome reconstruction from Bakta-annotated genomes revealed a small shared gene repertoire between SoF1 and H16. In the two-genome setting, pangenome frequency bins collapse such that “shell” corresponds to strain-specific clusters (present in only one genome; 50% frequency), whereas “soft-core” and intermediate frequency bins are not applicable. The pangenome comprised 9,804 genes in the CDS-related analysis, of which 925 were shared core genes (9.4%), 3,302 were SoF1-specific (33.7%), and 5,577 were H16-specific (56.9%) (Fig. 3B). Under the all-genes analysis, 9,814 genes were identified, including 1,118 shared core genes (11.4%), 3,189 SoF1-specific genes (32.5%), and 5,507 H16-specific genes (56.1%) (Fig. 3A). The slightly larger shared fraction in the all-genes view indicates that conserved features beyond CDS-defined loci also contribute to the shared genomic framework.

**Figure 3:**
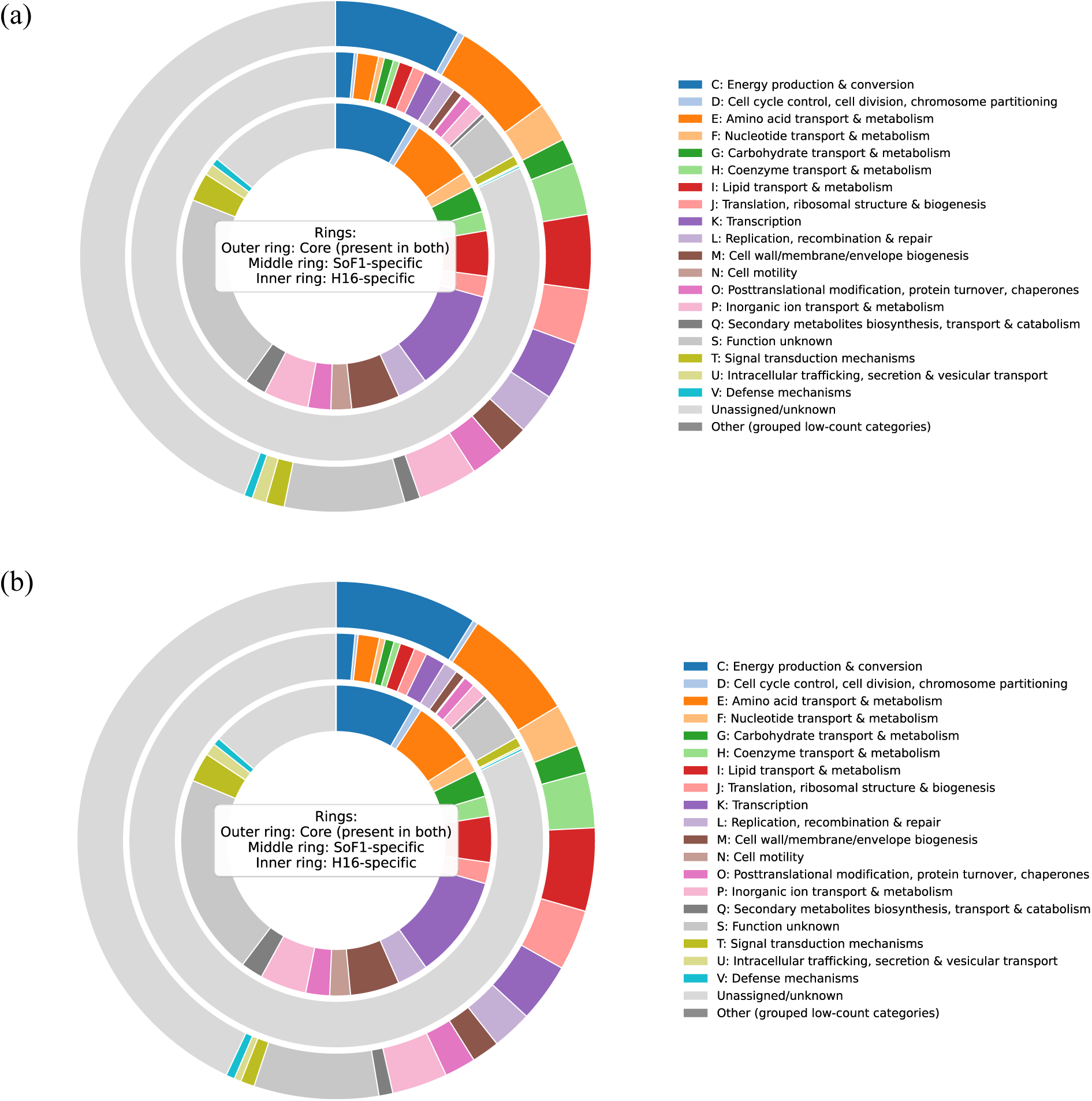
Pan-genome partitioning (**A**) all genes and (**B**) CDS-related between SoF1 and H16. PEPPAN pan-genome wheel generated from Bakta annotations including protein-coding and noncoding loci. Concentric rings depict the shared strict core (present in both genomes) and strain-specific gene sets (present in only SoF1 or only H16), colored by eggNOG/COG functional categories. The all-genes pan-genome comprises 9,814 clusters, partitioned into 1,118 core genes (11.4%), 3,189 SoF1-specific genes (32.5%), and 5,507 H16-specific genes (56.1%); “Unassigned/unknown” denotes genes without COG assignment. The CDS-related pan-genome comprises 9,804 clusters, including 925 core genes (9.4%), 3,302 SoF1-specific genes (33.7%), and 5,577 H16-specific genes (56.9%).

Functional partitioning of the strict core showed dominance of housekeeping and growth-supporting categories. Among annotated core genes, the most represented COG functional classes^28,29^ included Energy Production and Conversion (C), Amino acid Transport and Metabolism (E), and Lipid Transport and Metabolism (I) (Fig. **3**). The largest single core class remained Unassigned/unknown (all-genes: 494; CDS-related: 398), indicating a substantial conserved but poorly characterized fraction within the shared backbone.

### CO₂ fixation, H₂ oxidation, and Nitrogen metabolic gene contrasts

Across all reconstruction layers (KO/KEGG projection,^30^ Pathway Tools PGDBs,^31^ and ModelSEED^32^), both genomes encoded a conserved lithoautotrophic backbone consistent with aerobic growth on CO₂ and H₂. In the Pathway Tools reconstructions, PathoLogic inferred 436 MetaCyc pathways^33^ for H16 and 468 for SoF1, with 296 pathways shared (≈68% of H16 pathways and ≈63% of SoF1 pathways), indicating substantial conservation of core metabolism alongside meaningful chassis-specific specialization (Fig. **4A**). At the pathway-class level, both strains were dominated by biosynthesis and degradation/utilization functions, while SoF1 showed a higher number of biosynthetic pathways (178 vs 147) and energy/precursor pathways (36 vs 25), consistent with a broader inferred anabolic repertoire (Fig. **4B**).

**Figure 4:**
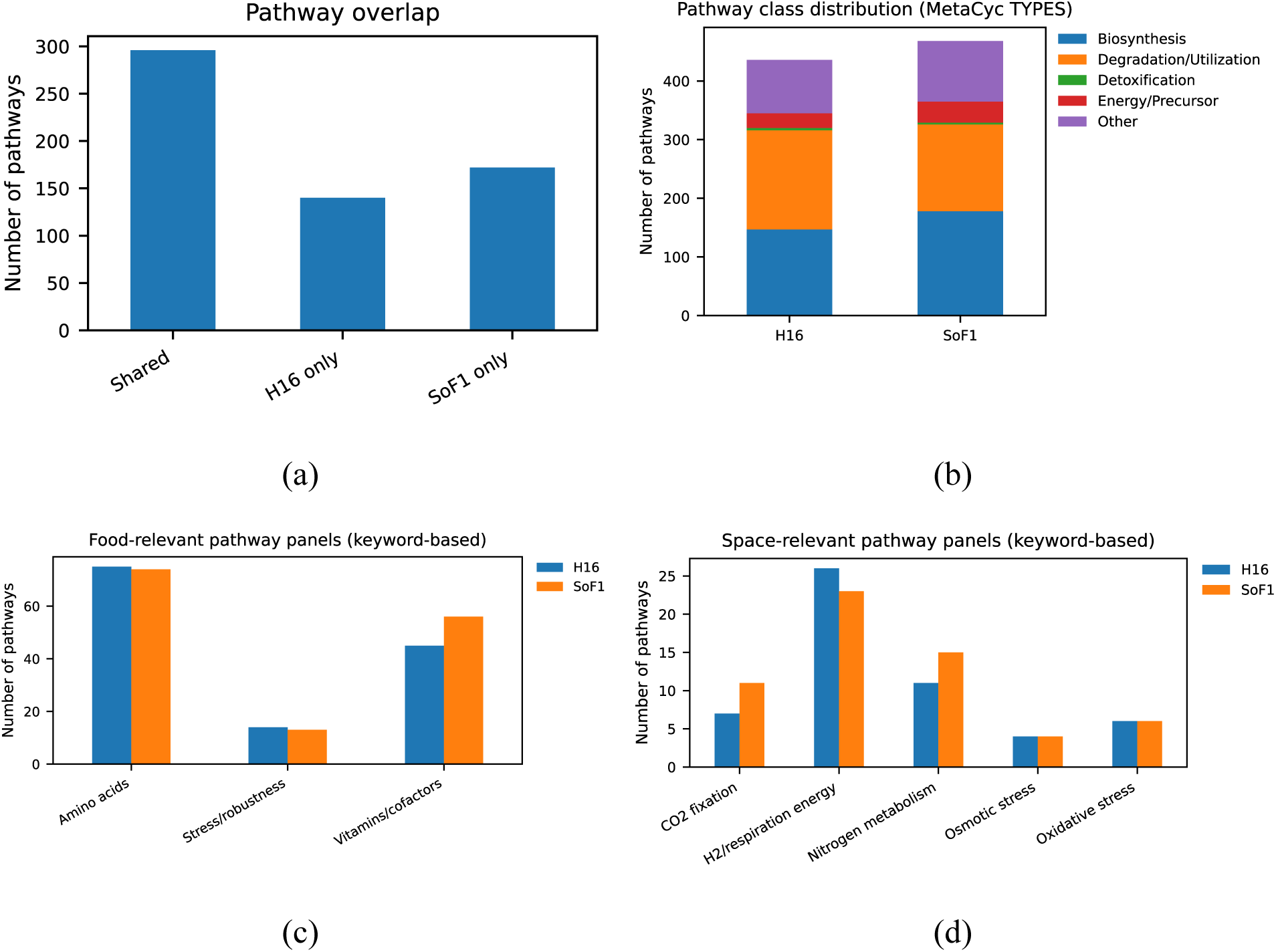
Pathway-level comparison of H16 and SoF1 PGDB reconstructions for gas-fermentation chassis selection. **A)** Pathway overlap between PathoLogic-inferred MetaCyc pathways in H16 and SoF1, showing shared and strain-specific pathway counts. **B)** Distribution of inferred pathways across top-level MetaCyc functional classes (Biosynthesis, Degradation/Utilization, Detoxification, Energy/Precursor, Other). **C)** “Food-relevant” pathway panel counts derived from PGDB exports, grouping pathways into amino-acid metabolism, vitamins/cofactors, and stress/robustness categories. **D)** “Space-relevant” pathway panel counts summarizing pathways associated with CO₂ fixation, H₂/respiration-linked energy metabolism, nitrogen metabolism, and oxidative/osmotic stress. Pathways were exported from each strain-specific PGDB and summarized computationally; panel assignments reflect targeted pathway grouping used for high-level functional comparison.

**Figure 5:**
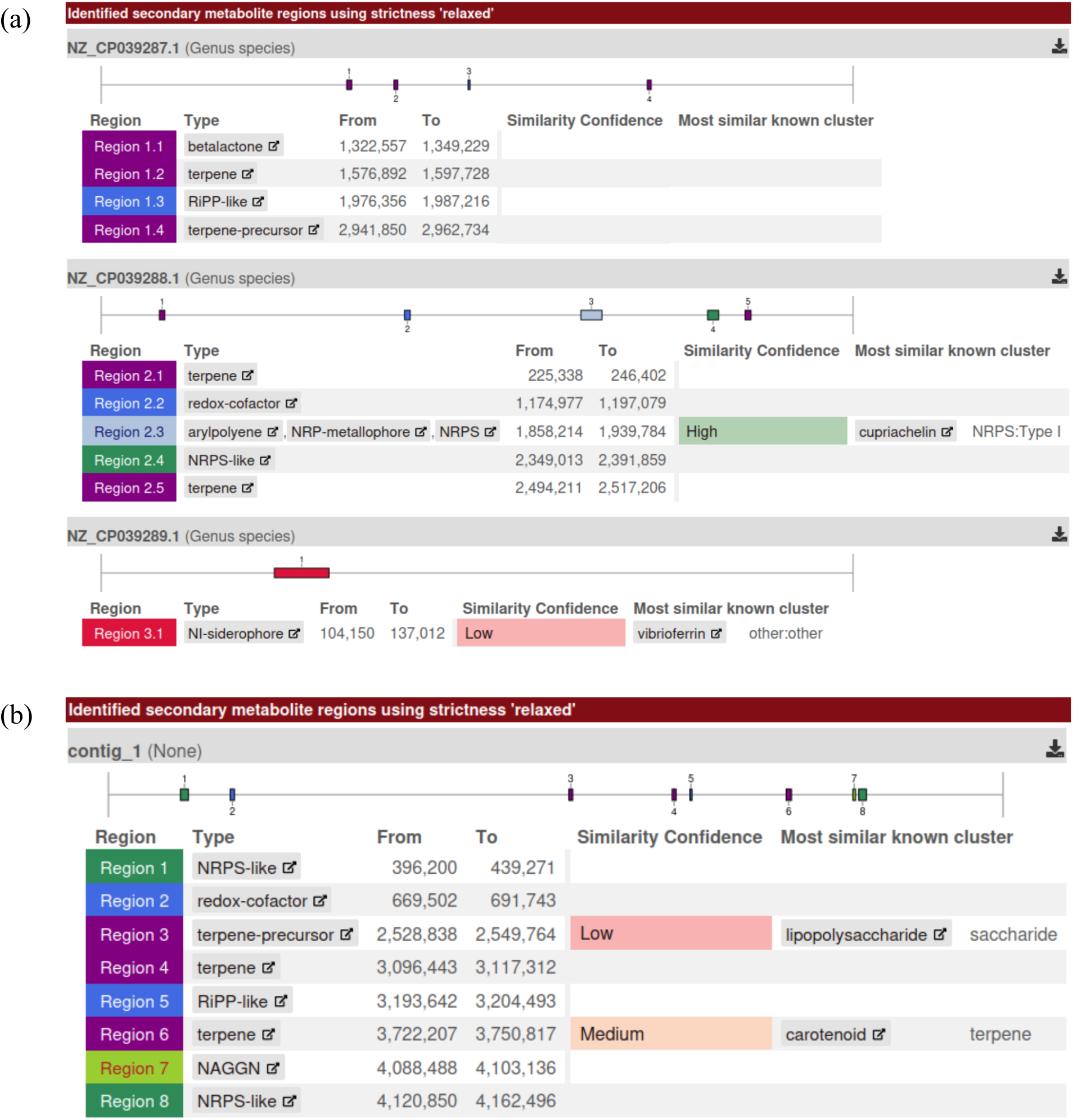
antiSMASH-predicted biosynthetic gene cluster repertoires in the two hydrogen-oxidizing food-protein chassis. **A**) *Cupriavidus necator* H16. **B**) *Xanthobacter sp.* SoF1. Maps show biosynthetic gene clusters (BGCs) predicted from the respective genome assemblies using antiSMASH (v7), with open reading frames rendered as arrows (direction indicates transcriptional orientation) and clusters annotated by predicted BGC class (e.g., NRPS-like, RiPP-like, terpene, redox-cofactor/PQQ biosynthesis, siderophore/metal-binding pathways, β-lactone, and hybrid NRPS–metallophore/arylpolyene where present). Cluster boundaries correspond to antiSMASH calls and should be interpreted as predicted specialized-metabolite loci rather than experimentally confirmed products.

Within this shared lithoautotrophic framework, CO₂ fixation capacity was supported by the presence of the CBB cycle in both PGDBs (CALVIN-PWY; Fig. **4D**), aligning with genome-encoded Rubisco/CBB modules and enabling “power-to-food” conversion of renewable H₂ and CO₂ into biomass. H₂ oxidation potential was similarly conserved at the gene level (hydrogenase and accessory maturation systems), supporting high-rate respiratory energy generation needed for rapid autotrophic growth.

Nitrogen acquisition encoded the strongest functional contrast. SoF1 uniquely carried a nitrogen-fixation pathway (N2FIX-PWY) and additional nitrogen-assimilation variants, consistent with N₂- supported growth predictions and DIAMOND-based confirmation of a complete *nifHDK* cluster^34^ with accessory/maturation genes in SoF1 and its absence in H16 (Fig. **4D**). Pathway inference further supported greater nitrogen-metabolism breadth in SoF1, including assimilatory nitrate-reduction potential, whereas H16 lacked the nitrogen-fixation program and instead relied on canonical ammonia assimilation routes (e.g., GS/GOGAT). Beyond nitrogen inputs, Pathway Tools highlighted additional carbon-flexibility differences, as H16 encoded the glyoxylate cycle (GLY-OXYLATE-BYPASS), which was absent in SoF1, consistent with greater potential to assimilate acetyl-CoA–derived C₂ substrates and to reutilize overflow acetate (Fig. 4A).^35^

Finally, both strains retained broad biosynthetic capacity relevant to food applications: amino-acid pathway coverage was comparable, while SoF1 showed more vitamin/cofactor-associated pathways in the PGDB-derived panel analysis (56 vs 45), which may reduce micronutrient supplementation needs in defined media (Fig. **4C**). Stress-relevant pathway counts (oxidative and osmotic stress) were similar between strains, supporting robustness under aerobic gas fermentation and downstream processing (Figs. **4C** & **4D**). Tables **3** and **4** summarize comparative presence and absence of CO_2_ fixation routes, hydrogenase class, and N_2_ acquisition pathways in both chassis.

**Table 3:** Comparison of carbon and hydrogen fixation pathways in SoF1 and H16.

| Functional module<br>(food/space relevance) | Key marker genes/<br>operon hallmark | SoF1 | H16 | Evidence / notes |
| --- | --- | --- | --- | --- |
| **CO<sub>2</sub> fixation class** |  |  |  |  |
| Primary CO<sub>2</sub> fixation<br>(CBB cycle) | *cbbL/cbbS* (RuBisCO),<br>*prk* (phosphoribuloki-<br>nase), *cbbR* (LysR reg-<br>ulator) | Yes | Yes<br>(dupli-<br>cated op-<br>erons) | H16 and SoF1 fixes<br>CO<sub>2</sub> via CBB; Ru-BisCO/other CBB<br>enzymes are in du-<br>plicate *cbb* operons<br>(one on pHG1, one<br>on chr2) with *cbbR*<br>at the chromosomal<br>locus.<sup>27,36</sup> |
| Anaplerotic CO<sub>2</sub> incor-<br>poration (flexibility on<br>mixed feeds / robust-<br>ness) | *ppc* (PEP carboxylase),<br>*pyc* (pyruvate carbox-<br>ylase) | *ppc* pre-<br>sent, *pyc*<br>absent | Yes | Common in many<br>aerobic proteobacte-<br>ria. |
| Formate oxidation / C1<br>redox (process flexibil-<br>ity; electro-/gas fermen-<br>tation hybrids) | *fds* (soluble NAD-re-<br>ducing FDH), mem-<br>brane-bound FDHs<br>(*fdh/fdo/fdw* clusters) | Yes | Yes (≥4<br>FDHs) | H16 encodes multi-<br>ple formate dehydro-<br>genases (one soluble<br>NAD-reducing +<br>several membrane-<br>associated sys-<br>tems).<sup>36</sup> |
| **Hydrogenase class (role in power-to-food)** |  |  |  |  |
| Membrane-bound up-<br>take [NiFe] hydro-<br>genase (energy-con-<br>serving H<sub>2</sub> oxidation) | Commonly *hupS/hupL* (+<br>membrane anchor/cyto-<br>chrome, plus *hyp* matura-<br>tion) | Yes | Yes | H16's hydrogenase re-<br>gion is concentrated on<br>pHG1 and includes the<br>uptake system. |
| Soluble NAD-reduc-<br>ing (bidirectional-<br>type) hydrogenase<br>(direct reducing<br>power for CO<sub>2</sub> fixa-<br>tion / biosynthesis) | H16-classically: *hoxF*-<br>*hoxU-hoxY-hoxH-hoxI* (+<br>*hyp* genes) | Yes | Yes | H16 SH operon and its<br>*hox/hyp* architecture are<br>described; high value<br>for H<sub>2</sub> technologies due<br>to activity and O<sub>2</sub> toler-<br>ance. |
| Regulatory (H<sub>2</sub>-sens-<br>ing) hydrogenase /<br>control system (sta-<br>bility in fluctuating<br>gas feeds) | Sensor hydrogenase<br>(*hoxB/hoxC* or<br>*hupU/hupV*) with two-<br>component regulators<br>(e.g., *hoxJ/hoxA*) | Yes | Yes | H16 uses a dedicated<br>regulatory hydrogen-<br>ase/control circuitry<br>governing hydrogenase<br>expression. |
| [NiFeSe] uptake hydrogenase (often high activity; potential O<sub>2</sub> tolerance depending on subgroup) | large + small subunits (selenocysteine-containing large subunit) | Yes | Not reported | SoF1 strain patent explicitly claims genes encoding NiFeSe hydrogenase large and small subunits (sequence listings). |

**Table 4:**
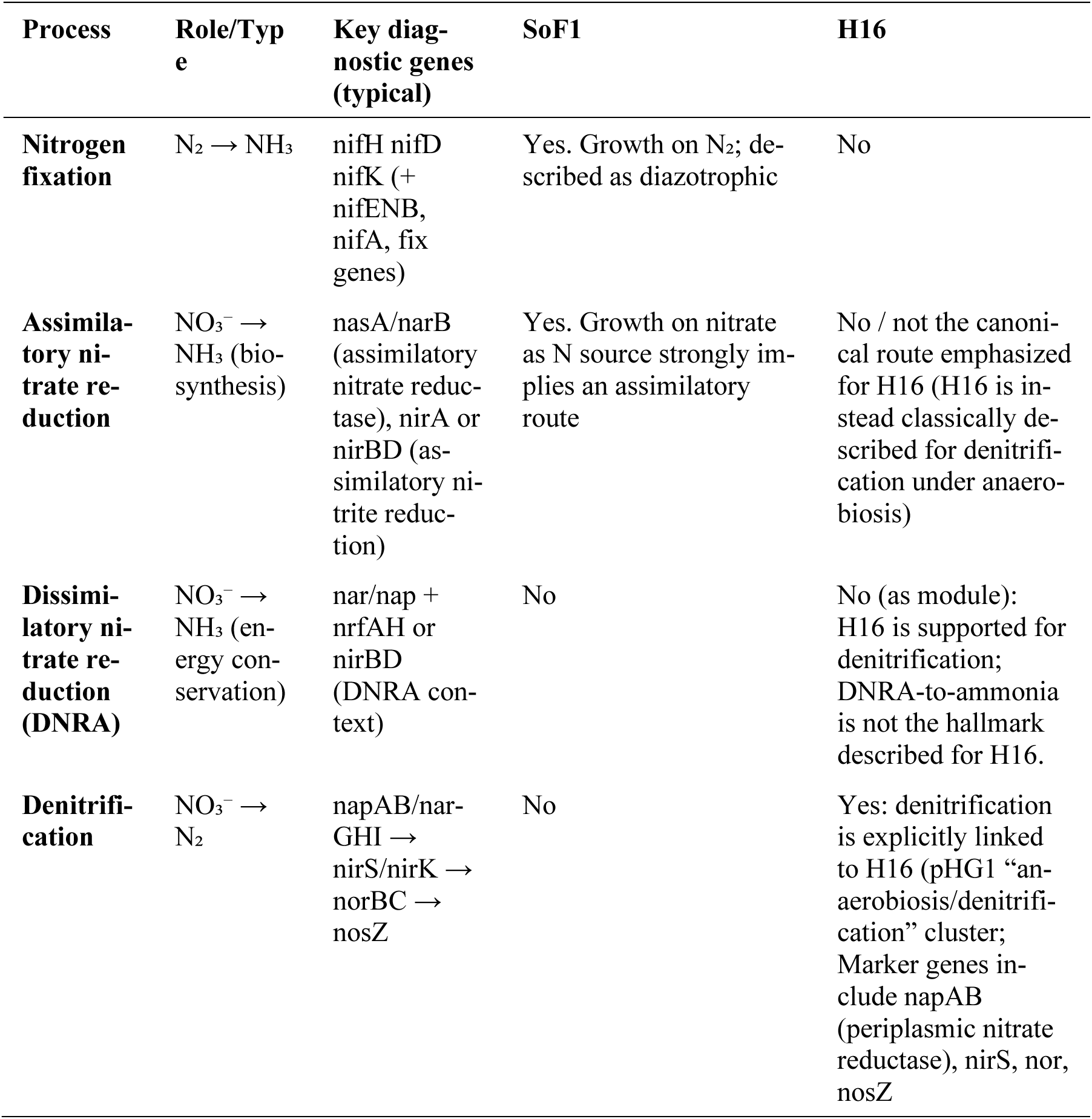
Comparison of nitrogen fixation pathways in SoF1 and H16.

| Process | Role/Type | Key diagnostic genes (typical) | SoF1 | H16 |
| --- | --- | --- | --- | --- |
| **Nitrogen fixation** | N<sub>2</sub> → NH<sub>3</sub> | nifH nifD<br>nifK (+<br>nifENB,<br>nifA, fix<br>genes) | Yes. Growth on N<sub>2</sub>; described as diazotrophic | No |
| **Assimilatory nitrate reduction** | NO<sub>3</sub><sup>-</sup> → NH<sub>3</sub> (biosynthesis) | nasA/narB (assimilatory nitrate reductase), nirA or nirBD (assimilatory nitrite reduction) | Yes. Growth on nitrate as N source strongly implies an assimilatory route | No / not the canonical route emphasized for H16 (H16 is instead classically described for denitrification under anaerobiosis) |
| **Dissimilatory nitrate reduction (DNRA)** | NO<sub>3</sub><sup>-</sup> → NH<sub>3</sub> (energy conservation) | nar/nap + nrfAH or nirBD (DNRA context) | No | No (as module): H16 is supported for denitrification; DNRA-to-ammonia is not the hallmark described for H16. |
| **Denitrification** | NO<sub>3</sub><sup>-</sup> → N<sub>2</sub> | napAB/narGHI → nirS/nirK → norBC → nosZ | No | Yes: denitrification is explicitly linked to H16 (pHG1 “anaerobiosis/denitrification” cluster; Marker genes include napAB (periplasmic nitrate reductase), nirS, nor, nosZ |

### Biosynthetic capacity for protein quality and stress- and process-relevant traits

Both genomes encode the core informational machinery expected of robust, fast-growing Proteobacteria (e.g. translation factors, aminoacyl-tRNA synthetases, and ribosomal proteins), consistent with high protein synthesis capacity under gas-fermentation regimes. Beyond housekeeping functions, genome-enabled process traits differed in their predicted surface and interaction repertoire (MacSyFinder/TXSScan). H16 carries a more extensive set of cell-surface and interaction systems, including flagellar genes, Tad and type IVa pili, a Type II secretion system, and a Type VI secretion system. By contrast, SoF1 shows a simpler surface-module profile, characterized mainly by Type I secretion, multiple Type Va autotransporters, and a single pT4SSt-like module, while lacking predicted flagellar and T6SS machinery. These differences are likely to influence process-relevant traits such as cell aggregation, surface attachment, shear sensitivity, and broth clarification, and therefore highlight practical targets for chassis streamlining.

antiSMASH profiling suggests that both strains devote only a modest fraction of their genomes to secondary metabolism, and that this chemistry is oriented toward environmental fitness rather than toxin production. The predicted repertoires are not dominated by polyketide synthase (PKS) or non-ribosomal peptide synthetase (NPRS) cytotoxin architectures commonly associated with pathogenic metabolite risk. SoF1 encodes eight biosynthetic gene cluster (BGC) regions, including NPRS-like, terpene, ribosomally synthesized and post-translationally modified peptide (RiPP)-like, pyrroloquinoline quinone (PQQ), and a strain-specific N-acetylglutaminylglutamine amide (NAGGN) locus linked to osmoprotection. H16 encodes ten BGC regions across its three replicons, including beta-lactone, terpene, RiPP-like, PQQ, NPRS-like, and siderophore or metallophore-associated clusters. Together, these patterns are more consistent with stress adaptation, competition, and metal homeostasis than with toxin-class metabolite production. These differences are engineerable in the sense that they identify specific loci that can be preserved to enhance process robustness or removed to streamline the chassis and minimize unnecessary secondary-metabolite potential in food-grade applications.

### Safety-relevant genomic scan

To assess food-grade suitability, we screened for genomic features that are directly relevant to biosafety evaluation: antimicrobial resistance (AMR) determinants, their potential mobility through plasmids or integrons, virulence-associated genes and secretion systems, and matches to large toxin-reference databases. These categories are important because they address the main regulatory concerns for production strains namely transferable resistance, pathogenicity-associated functions, and toxin-encoding potential. Under the strict thresholds used, both chassis displayed a low-risk genetic profile.

#### AMR (RGI/CARD + AMRFinderPlus)

AMR genes are safety-relevant primarily when they represent acquired determinants or are associated with genetic contexts that could enable dissemination to other microorganisms. Following EFSA guidance for WGS-based characterisation of microorganisms intentionally used in the food chain, AMR- and virulence-associated homologs were reported at ≥80% sequence identity and ≥70% subject-length coverage.^37,38^ To distinguish close, high-confidence matches from distant functional homologs, we additionally flagged a conservative subset using ∼≥90% identity (and high-length coverage), consistent with default BLAST-based calling behaviour in AMRFinderPlus/NCBI pathogen-detection workflows.^39–41^ Under this high-stringency filter, no AMR determinants met the cutoff in either genome (H16: RGI 0 at ≥90% identity; AMRFinderPlus 0 at ≥90%; SoF1: RGI 0 at ≥90%). At the EFSA reporting threshold, H16 retained only a few AMR-related hits consistent with intrinsic background functions, such as efflux-associated homologs. These were considered review items rather than evidence that H16 carries acquired, clinically relevant AMR genes.

#### Mobility and integrons

Mobility-associated elements such as plasmids, transposon-linked regions, and integrons are important because they increase the likelihood that resistance or other unwanted traits could be horizontally transferred. IntegronFinder reports no complete integrons, CALIN, or In0 elements in either genome. MOB-suite indicates minimal plasmid-like architecture overall (H16: 3 contigs with 1 plasmid-like; SoF1: 1 contig with 0 plasmid-like), and the combined evidence does not support integron-mediated cassette accumulation or obvious mobile AMR islands. *Virulence-factor homology (ABRicate/VFDB)*: Virulence-related genes and secretion systems are safety-relevant because they can indicate host interaction, colonization, or pathogenicity-associated functions, although conserved environmental systems must be interpreted cautiously. Under strict filtering (≥90% identity and ≥80% coverage), zero VFDB hits were retained for both strains. At lenient thresholds (≥75% identity, ≥50% coverage), H16 yields 18 hits and SoF1 yields 2 hits, dominated by generic functions (motility/chemotaxis, pili/biofilm-associated components, secretion-associated homologs) rather than toxins or hallmark invasion/colonization factors. Critically, no T3SS is predicted by MacSyFinder in either strain, which strongly reduces the likelihood of classical Gram-negative pathogenicity island architecture.

#### Large toxin database scan (DIAMOND vs DBETH + Toxinome)

Large toxin-database searches are included to identify canonical toxin families or close toxin-like homologs that could represent an exclusion criterion or require targeted follow-up during strain qualification. Under strict DIAMOND thresholds (id ≥90%, qcov ≥80%, e ≤1e-20), no canonical food-borne toxins were detected for both strains (H16: 0; SoF1: 0). In H16, two proteins fall into toxin-like families requiring context review (RTX-like and Tc-like), but their genome context does not show integron linkage or mobility-keyword enrichment in the neighborhood, and the RTX-like hit lacks supportive signals for a canonical RTX secretion/activation module in the immediate vicinity. SoF1s toxinome matches are limited to generic toxin-like annotations (e.g., AAA+ family hits) rather than canonical exotoxins and are not supported by hallmark virulence system context.

Overall, the independent safety screens covering antimicrobial resistance, mobility and integrons, virulence-factor homology, secretion systems, and two toxin databases did not detect high-confidence foodborne exotoxins in either genome under strict criteria. Only a few context-review items remained, most notably RTX- and toxin complex-like annotations in H16, which require targeted domain verification and locus-level inspection rather than indicating confirmed toxin risk.

### Actionable implications for sustainable protein platforms

From a production and qualification standpoint, the genome-informed screens suggest that both chassis can be advanced with a risk-managed, evidence-first qualification plan rather than requiring broad “de-risking” redesign. The BGC comparison indicates that SoF1’s secondary metabolism is weighted toward terpenes/precursors and a distinctive NAGGN osmoadaptation locus, aligning with hypotheses around robustness under gas-fermentation and drying stresses. H16 carries additional and more distributed biosynthetic diversity, including metal acquisition chemistry (metallophore/siderophore-like regions, including a megaplasmid-encoded NI-siderophore) and a β-lactone region, motivating targeted analytics to ensure that non-essential secondary products are not expressed in a way that could complicate food qualification or sensory outcomes.

On safety, the compiled outputs support a narrative aligned with EFSA/FDA-style concerns: no integrons, no high-confidence acquired AMR under stringent identity thresholds, no T3SS/T4bP, and no canonical foodborne exotoxins detected by strict DIAMOND screening.^38^ At the same time, the results argue for transparent disclosure of the few context-review flags (mobility machinery present without AMR; H16’s single T6SSi instance; RTX/Tc-like hits requiring domain/synteny confirmation). This positions the chassis as non-pathogenic, industrially tractable Proteobacteria whose remaining “interaction biology” can be monitored and minimized (if needed) through targeted strain engineering once expression under production conditions is empirically established.

## Discussion

HOB offer a credible route to “protein without farms” or land-free biotechnology because their core metabolism converts CO₂ into biomass using reducing power from H₂, allowing production that is largely independent of arable land, climate variability, and fertilizer-driven nitrogen disruption.^6,10^ Our comparative genomics of H16 and SoF1 shows that this promise rests on a conserved autotrophic backbone, yet the chassis that carry it are deeply different in genome architecture, accessory gene space, and nutrient economy differences that matter for industrial control, compositional consistency, and food-grade qualification. This distinction is central to the power-to-food thesis, i.e. platform choice cannot be justified by the presence of CO₂ fixation and hydrogenases alone but should be grounded in genome-resolved predictions of robustness, controllability, and the size of the “unknown” functional space that remains to be de-risked.

The strains’ architecture implies different engineering and regulatory narratives. H16’s multipartite genome distributes substantial accessory capacity across chromosome 2 and the mega-plasmid pHG1, reinforcing a modular organization in which major traits can be encoded on secondary replicons. Modularity can accelerate engineering by enabling targeted edits to functionally dense regions, but it also raises stability questions that regulatory bodies will reasonably prioritize for food applications, where consistent identity across passages and minimal mobile-risk features are critical.^27^ By contrast, SoF1’s single-replicon organization and compact genome support a streamlined chassis narrative that may simplify identity-by-sequence control and stability documentation. However, a compact genome does not imply complete functional understanding. Even model organisms retain sizable uncharacterized fractions (e.g., ∼15-20% in *E. coli* K-12 and ∼21% in *B. subtilis*), so the ‘function unknown’ component observed in both H16 and SoF1, including within the shared backbone, means genome-based feature confirmation should be paired with targeted functional validation and gene-product-focused analytics.^42–44^

At the metabolic level, both strains encode the canonical power-to-food machinery, e.g. a complete CBB cycle for CO₂ fixation and multiple [NiFe]-hydrogenase systems supporting H₂ oxidation, consistent with lithoautotrophic gas fermentation. The comparative value lies in how each chassis supplies, regulates, and buffers these functions under operational variability. H16 encodes duplicated carbon-fixation operons together with a more differentiated hydrogenase architecture, suggesting greater regulatory redundancy and metabolic flexibility during autotrophic growth. Rather than implying deliberate variation in feed-gas composition, these features may be especially relevant under realistic bioprocess heterogeneities, such as local gradients in hydrogen, oxygen, or redox state within the reactor. By contrast, SoF1 appears to implement the same core hydrogen-oxidizing autotrophic metabolism in a more compact genome, with fewer parallel modules but clear relevance as a production strain for edible biomass. These differences point to distinct engineering logics: H16 provides a more elaborate regulatory and metabolic framework that may better buffer fluctuating intracellular or reactor-scale conditions, whereas SoF1 represents a genetically more compact production chassis that may be advantageous for simplification and targeted optimization.

Nitrogen metabolism is one of the clearest functional differences between H16 and SoF1, and this difference becomes especially important under process conditions where nitrogen supply is limiting or costly. Comparative genomics indicates that SoF1 encodes both nitrogen fixation and nitrate assimilation, suggesting broader flexibility in how it can obtain nitrogen, including from atmospheric N₂ and oxidized nitrogen sources such as nitrate. By contrast, H16 lacks the genetic basis for nitrogen fixation and therefore depends more strongly on externally supplied fixed nitrogen in the medium, such as ammonia, ammonium salts, or potentially urea after hydrolysis. This distinction is not simply a matter of media formulation; it affects which chassis is better suited to industrial settings. In terrestrial biomanufacturing, a strain with broader nitrogen-use flexibility may reduce dependence on concentrated reduced-nitrogen inputs and expand medium design options. In closed-loop or highly resource-constrained systems, including space life-support concepts, the ability to access nitrogen from N₂ or nitrate could simplify nutrient recycling and reduce the need to store or continuously supply fixed nitrogen sources such as ammonium. The genomic comparison therefore identifies nitrogen metabolism as an important design parameter for selecting and engineering HOB chassis for different deployment scenarios.

Yield predictability depends not only on the ability to make biomass but also on how carbon is partitioned and how the cells tolerate process stress. Both genomes encode complete translational machinery and multiple rRNA operons, supporting high biosynthetic capacity. Yet both also encode PHB biosynthesis potential, implying that carbon diversion into storage polymers is a native routing option in the wild type. For a protein-focused process, this reinforces PHB attenuation (or dynamic control) as a rational, chassis-agnostic engineering lever to stabilize protein-rich biomass output and reduce biomass composition drift across cultivation states. Stress tolerance is similarly central to translation, i.e., both strains encode robust oxidative stress defenses and conserved heat-shock/chaperone systems that are relevant to high-aeration gas fermentation and downstream stabilization. SoF1’s enrichment for desiccation/DNA-protection features is consistent with improved tolerance to processing stresses such as osmotic transitions or drying, but the appropriate posture is to treat this as a genomics-derived hypothesis that should be validated under industrially realistic perturbations (gas-transfer limitation, oxidative load, osmotic challenge, and drying conditions) using defined metrics for productivity and composition.

Secondary metabolism highlights a productive tension between robustness and food-grade simplicity. H16’s broader BGCs repertoire including siderophore/metallophore and β-lactone-associated clusters may confer competitive fitness and stress protection, potentially enhancing culture resilience under nutrient limitation or oxidative pressure. At the same time, expanded secondary metabolism increases the metabolite space that may require monitoring or mitigation to satisfy compositional-consistency and safety expectations. The key interpretive point is not that either strain is “unsafe,” but that genome-resolved BGC landscapes should be treated as a structured riskand-opportunity register such that they indicate which metabolites might appear under specific physiological states and therefore where process control, analytical monitoring, or genetic attenuation can simplify regulatory narratives. In this context, motif-based transcription factor binding site (TFBS) predictions are useful for generating hypotheses about conditional induction, but mechanistic claims should be restrained unless supported by expression and metabolite data. A defensible translational strategy could be to explicitly test worst-case induction regimes (oxygen limitation, late stationary phase, nutrient stress), measure metabolite outputs, and delete or silence high-priority clusters where a “cleaner” food-grade chassis is desired.

Safety-oriented genome screening provides strong negative evidence that both strains lack canonical virulence determinants, major toxin families, and high-confidence transferable AMR architectures under stringent thresholds, aligning with baseline expectations for environmental Gram-negative production organisms. These results strengthen the safety-by-design foundation, but they do not replace empirical testing particularly given the persistent fraction of uncharacterized genes and the presence of specialized metabolism loci. The most persuasive pathway to acceptance therefore combines genome-based identity and hazard screening with targeted functional validation: stability testing across passages, analytics to confirm the absence (or tight control) of bioactive products under manufacturing conditions, and process specifications that manage predictable Gram-negative features such as LPS through downstream processing and quality control. Both genomes (H16 and SoF1) support strong potential as food-grade HOB platforms. SoF1 currently has more publicly visible food-facing safety documentation, while H16 has deeper mechanistic and engineering maturity; aligning both chassis under comparable safety and compositional validation would enable an equivalently evidence-based readiness assessment.

Taken together, as depicted in Fig. 6, comparative genomics reveals that that the here investigated “power-to-food” microbes share a conserved biochemical engine but differ profoundly in the genomic design choices that determine whether that engine can be operated predictably, efficiently, and in a manner consistent with food-grade regulation. SoF1’s compact genome and nitrogen-flexible potential strengthen its case as a deployable production host, particularly where nutrient inputs and closed-loop operation are central constraints. H16’s modular, replicon-parti-tioned architecture and regulatory-rich hydrogenotrophic machinery position it as a powerful chassis for mechanistic control, robustness engineering, and transferable design principles. More broadly, our analysis establishes a genomics-first framework for selecting and engineering HOB platforms: define the shared autotrophic minimum set, quantify and interpret the accessory space that drives operational phenotype, and couple hazard-aware genome screening to targeted validation and rational edits (e.g., PHB attenuation, selective BGC deletions, stability-focused modifications). This integration provides a realistic path toward scalable protein production without farms on Earth and a blueprint for closed-loop protein synthesis in space systems powered by renewable electricity.

**Figure 6:**
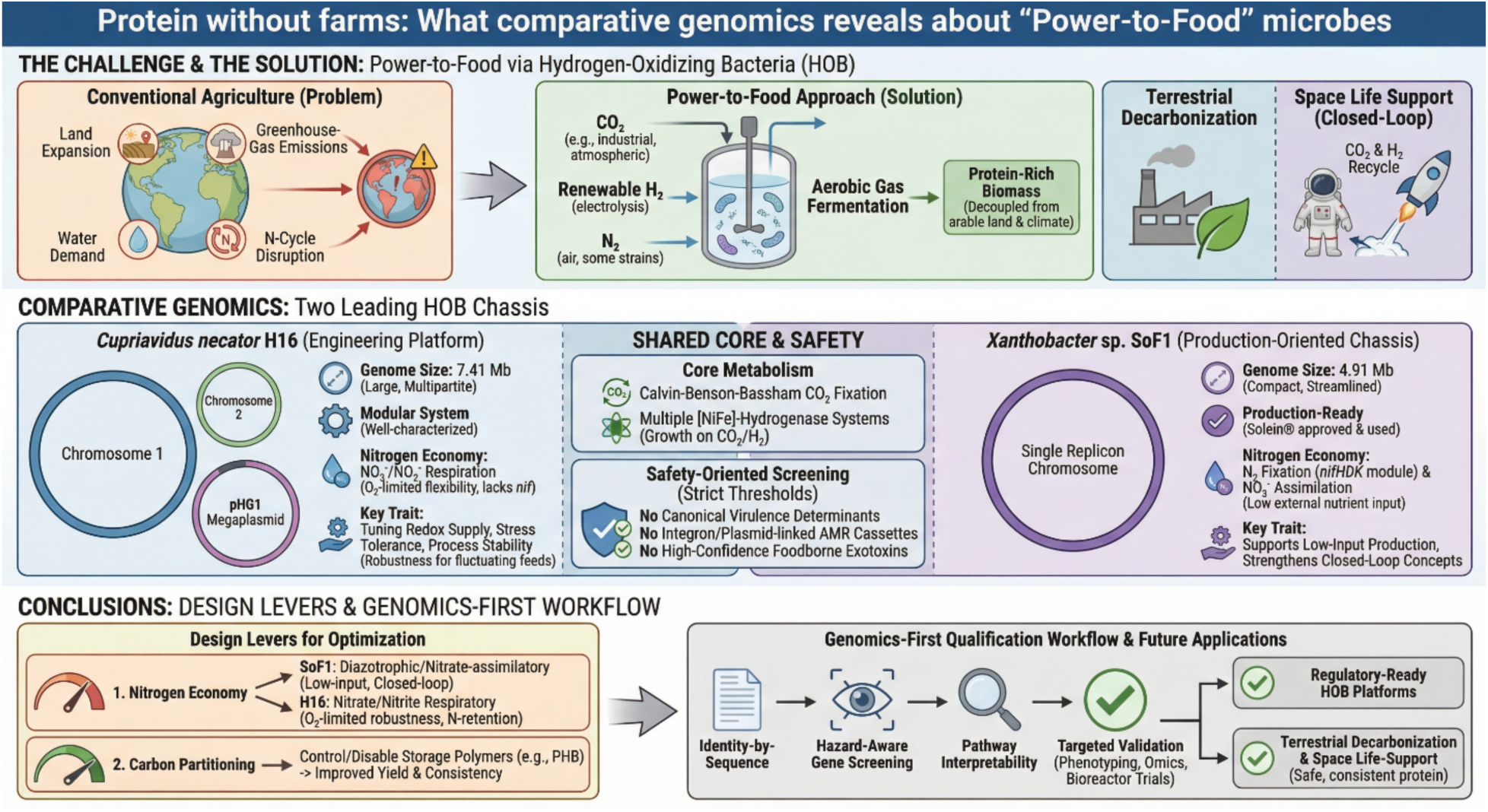
Comparative genomics highlights shared autotrophic core metabolism but distinct design-relevant traits in *Cupriavidus necator* H16 and *Xanthobacter* sp. SoF1, positioning hydrogen-oxidizing bacteria as promising power-to-food chassis for terrestrial decarbonization and closed-loop space biomanufacturing.

## Conclusions

Comparative genomics identifies SoF1 as the more streamlined, production-oriented HOB chassis for food applications, with a compact single-replicon genome and nitrogen-acquisition capacity that can reduce external nutrient inputs under constrained operation. Independently of genomics, SoF1 is already being used as the production organism for Solein®^45^ and has received regulatory approval for sale in Singapore. It has also been featured in consumer-facing products in that market, consistent with active commercialization rather than a purely prospective translation claim.^20,46^ At the same time, the genome architecture and regulatory logic revealed here highlight why H16 remains strategically valuable for advancing power-to-protein as an engineered platform. Its larger, multipartite genome and the historical concentration of hydrogenotrophic machinery on the mega plasmid pHG1 provides a well-characterized, modular system for dissecting and tuning redox supply, stress tolerance, and process stability traits that determine whether HOB bioprocesses can run reliably across fluctuating gas feeds and industrial duty cycles. Beyond host selection, our results point to concrete next steps for the field. Nitrogen economy is a primary design lever: SoF1’s diazotrophic and nitrate-assimilatory potential supports low-input production and strengthens closed-loop concepts, while H16’s nitrate/nitrite respiratory capacity motivates strategies for oxygen-limited robustness and nitrogen retention where fixed nitrogen is supplied.^10^ Carbon partitioning is similarly engineerable: disabling or tightly controlling storage-polymer routes (e.g., PHB) is a direct path toward improved yield predictability and compositional consistency.^21,47^ Finally, the genomics-first qualification workflow established here is reusable for screening new HOB candidates by integrating identity-by-sequence, hazard-aware gene screening, and pathway interpretability to support regulatory dossiers. Coupled with targeted validation (growth phenotyping, proteomics, metabolomics, and controlled bioreactor trials), this framework can accelerate regulatory-ready HOB platforms for terrestrial decarbonization and for space life-support, where recycling CO₂ and minerals into safe, consistent protein is mission-critical.

## Methods

The overview of computational pipeline and steps followed in the present study for genome comparison of two leading HOB chassis strains, *Cupriavidus necator* H16 and *Xanthobacter* sp. SoF1 is shown in Fig. 7. Detailed procedure in a step-by-step manner followed in this study is presented in following subsections.

**Figure 7:**
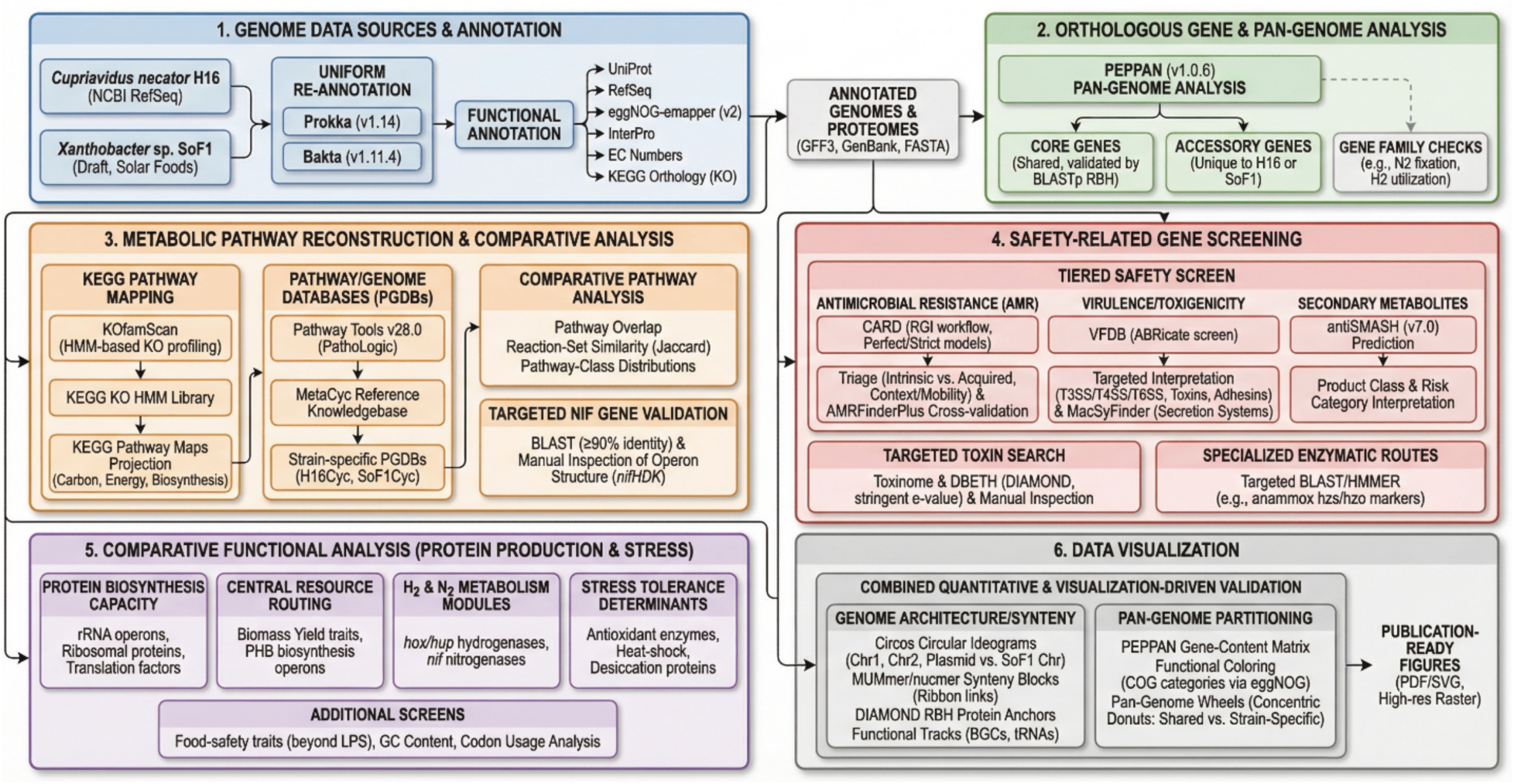
Overview of the comparative genomics workflow used to annotate, compare, and safetyscreen *Cupriavidus necator* H16 and *Xanthobacter* sp. SoF1, integrating pan-genome analysis, metabolic reconstruction, targeted validation, and visualization.

### Genome data sources and annotation

The complete genome sequence of *Cupriavidus necator* H16^18^ was obtained from NCBI RefSeq^48^ (accession number/identifier in Table **1**), and the draft genome of *Xanthobacter sp.* SoF1^21^ was provided by Solar Foods Oyj^20^ (Vantaa, Finland). The SoF1 genome, originally assembled from shortand long-read sequencing data, was received as assembled 1 contig. To ensure uniform annotation, both genomes were re-annotated using Prokka (v1.14).^49^ Prokka was run with default parameters to predict protein coding sequences (CDS), tRNAs, rRNAs, and other features, producing standard GenBank-format outputs. In parallel, the Bakta pipeline (v1.11.4) was employed for a second annotation of each genome, leveraging its alignment-free gene identification for rapid, standardized results.^50^ Annotation outputs from Prokka and Bakta were compared to reconcile any discrepancies in gene calls and functional assignments. Predicted proteins were functionally annotated by homology to UniProt,^12^ RefSeq,^48^ eggNOG-emapper (v2),^51^ and InterPro^52^ databases, and assigned Enzyme Commission (EC) numbers^53^ and KEGG Orthology (KO)^30^ identifiers when possible.

### Orthologous Gene and Pan-Genome Analysis

We performed a two-genome pan-genome analysis using PEPPAN (v1.0.6).^54^ PEPPAN was provided with bakta-generated GFF3 annotation files from both genomes and clustered genes based on sequence similarity (minimum 95% BLASTp identity). This yielded a pangenome matrix distinguishing core genes present in both genomes from accessory genes unique to each. Orthologous gene pairs were further validated by bidirectional best BLAST hits. Gene families of interest, such as those related to nitrogen fixation or hydrogen utilization, were specifically checked for presence/absence in each genome.

### Metabolic Pathway Reconstruction and Comparative Analysis

We reconstructed and compared metabolic pathways using a multi-layer strategy that enables cross-validation of pathway calls across independent annotation frameworks. First, predicted proteins from uniform genome annotations were assigned KEGG Orthology (KO)^55^ identifiers using KOfamScan (HMM-based KO profiling) against the KEGG KO HMM library. KO assignments were then projected onto KEGG pathway maps to provide a genome-wide overview of central carbon metabolism, energy conservation, and biosynthetic capacity (e.g., amino acid, vitamin/cofactor, and lipid metabolism). In parallel, we built strain-specific Pathway/Genome Databases (PGDBs)^56^ for H16 and SoF1 using Pathway Tools v28.0 (PathoLogic)^31^ with MetaCyc^33^ as the reference knowledge base. GenBank files generated from Prokka/Bakta were imported into Pathway Tools, and PathoLogic was executed in batch mode to infer pathways from enzyme and geneproduct evidence. The resulting PGDBs (H16Cyc and SoF1Cyc) enabled direct strain-to-strain comparison of inferred pathway content. To support fully reproducible, GUI-free analyses, pathway tables were programmatically exported from each PGDB using a custom Pathway Tools Lisp routine that extracts pathway identifiers, names, reaction membership, gene membership, and MetaCyc pathway class assignments (TYPES) retrieved from the MetaCyc template frames via the INSTANCE-OF relationship. Exported tables were parsed in Python to compute pathway overlap (shared vs strain-specific pathways), reaction-set similarity (Jaccard index) for shared pathways, and pathway-class distributions; results were written to CSV/Excel and visualized as publication-quality vector (PDF/SVG) and high-resolution raster figures. To validate nitrogen-fixation predictions, we performed targeted BLAST searches of both genomes against curated *nif* gene sequences using a ≥90% amino-acid identity threshold, followed by manual inspection of genomic neighborhoods to confirm operon structure and the presence of key accessory/maturation genes (e.g., *nifHDK* core and associated assembly/regulatory components).

### Safety-Related Gene Screening

To assess food-grade suitability and anticipate regulatory concerns, we performed a tiered, genome-informed safety screen focused on (i) antimicrobial resistance (AMR),^57^ (ii) virulence/toxigenicity, and (iii) undesired secondary metabolite potential. AMR determinants were queried using the Comprehensive Antibiotic Resistance Database (CARD)^58^ via the Resistance Gene Identifier (RGI) workflow (local database; DIAMOND-based alignments), prioritizing high-confidence calls under surveillance-style stringency (Perfect/Strict models and/or ≥90% sequence identity with ≥90% query coverage). Hits were further triaged as likely intrinsic versus acquired, clinically relevant resistance by inspecting gene context and mobility signals, including proximity to transposases/integrases and association with plasmid/integron features (IntegronFinder; MOB-suite), and by cross-validating calls with AMRFinderPlus for concordant gene family assignments. Putative virulence factors were screened by homology against the Virulence Factor Database (VFDB),^59^ using an automated genome screen (ABRicate/VFDB) followed by targeted interpretation that emphasized hallmark pathogenicity systems secretion system islands (T3SS/T4SS/T6SS), toxins, adhesins, and invasins rather than ubiquitous Gram-negative envelope or housekeeping components. Presence/absence of secretion-system modules was additionally assessed using a dedicated detection framework (MacSyFinder/TXSScan models) to minimize false positives from partial homologs and to distinguish complete systems from isolated components. Secondary metabolite biosynthetic gene clusters were predicted using antiSMASH (v7.0)^60^ on uniformly annotated GenBank inputs, and detected clusters were interpreted by predicted product class and known risk categories (e.g., toxin-associated metabolites versus iron-scavenging siderophores). To complement database-driven calls, we conducted targeted homology searches against curated toxin reference sets (hemolysins, enterotoxins, neurotoxins) from Toxinome^61^ and DBETH^62^ databases using DIAMOND with stringent e-value and coverage cutoffs, followed by manual inspection to exclude spurious low-complexity matches (SEG filtering) and fragmentary alignments. Finally, we explicitly screened for particularly concerning enzymatic routes reported in specialized taxa (e.g., anammox-associated hydrazine machinery such as *hzs/hzo* markers) using targeted BLAST/HMMER searches against curated reference sequences to confirm the absence of such pathways.

### Comparative Functional Analysis (Protein Production and Stress Tolerance)

To evaluate suitability as single-cell protein chassis, we performed a targeted comparative genomics analysis of features linked to (i) protein biosynthesis capacity (rRNA operon copy number, ribosomal proteins, translation factors), (ii) central resource-routing traits affecting biomass yield (e.g., PHB biosynthesis operons^21^), (iii) hydrogen and nitrogen metabolism modules supporting autotrophic growth (screening for *hox*/*hup* hydrogenase^63^ clusters and *nif* nitrogenase^34^ genes), and (iv) stress tolerance determinants relevant to cultivation and downstream processing (antioxidant enzymes, heat-shock systems, desiccation-associated proteins). We additionally screened both genomes for potentially problematic food-safety traits (e.g., toxin-associated features beyond baseline Gram-negative LPS) and assessed GC content and codon usage as indicators of atypical genomic composition that could confound expression or processing.

### Data Visualization

To aid interpretation of the comparative genomics results, we combined quantitative outputs with visualization-driven validation focused on (i) genome architecture/synteny and (ii) pan-genome partitioning with functional context. For genome architecture, the complete multipartite H16 genome (Chr1, Chr2, and megaplasmid) and the SoF1 chromosome were visualized as circular ideograms in Circos^26^ using custom karyotype files. Synteny blocks were computed from wholegenome alignments generated with MUMmer/nucmer^64^ and plotted as ribbon links to highlight conserved segments and large-scale rearrangements. To reinforce nucleotide-level synteny with gene-level correspondence, we additionally derived protein anchors via DIAMOND^65^ blastp reciprocal best hits (RBH) from Bakta-predicted proteomes and overlaid these as high-confidence orthology links. Functional feature tracks were added to contextualize structural conservation: biosynthetic gene cluster (BGC) coordinates were extracted from antiSMASH^60^ GenBank outputs and plotted as highlight tracks, and tRNA loci were included as an auxiliary genomic feature layer. For pan-genome structure, the PEPPAN gene-content matrix (Rtab) built from Bakta annotations was used to classify each pan-gene as shared core (present in both genomes) or strain-specific (SoF1-only vs H16-only). Functional coloring of pan-genes was assigned by mapping representative locus tags to eggNOG^51^/COG categories^28^ from the processed annotation tables; genes without COG calls were labeled Unassigned/unknown, and sparse categories were collapsed into an Other bin (minimum count = 10). Pan-genome wheels were rendered as concentric donut rings separating shared versus strain-specific gene space and exported as publication-ready PNG (600 dpi) and SVG via a command-line Python plotting workflow. All plotting scripts, configuration files, and intermediate matrices were versioned and deposited alongside the analysis outputs to ensure full reproducibility.

## Abbreviations

antiSMASH: antibiotics & Secondary Metabolite Analysis SHell
BGCs: Biosynthetic Gene Clusters
BLAST: Basic Local Alignment Search Tool
CARD: Comprehensive Antibiotic Resistance Database
CBB: Calvin–Benson–Bassham (cycle)
CDS: Coding Sequences
COG: Clusters of Orthologous Genes
CRISPR: Clustered Regularly Interspaced Short Palindromic Repeats
DIAMOND: Double Index Alignment of Next-generation sequencing Data sequence aligner
EFSA: European Food Safety Authority
FDA: U.S. Food and Drug Administration
G+C: Guanine plus Cytosine (GC content)
GRAS: Generally Recognized As Safe
H16: *Cupriavidus necator* H16 (strain designation)
HGT: Horizontal Gene Transfer
HOB: Hydrogen-Oxidizing Bacteria
KEGG: Kyoto Encyclopedia of Genes and Genomes
KO: KEGG Orthology
MetaCyc: MetaCyc Metabolic Pathway Database
NADH: Reduced Nicotinamide Adenine Dinucleotide
NADPH: Reduced Nicotinamide Adenine Dinucleotide Phosphate
NAGGN: N-acetylglutaminylglutamine amide
NCBI: National Center for Biotechnology Information
NRPS: Nonribosomal Peptide Synthetase
OG: Orthologous Groups
PEPPAN: Phylogeny Enhanced Pipeline for PAN-genome
PGDB: Pathway/Genome Database
PHB: Poly(3-hydroxybutyrate) (polyhydroxybutyrate)
PQQ: Pyrroloquinoline Quinone
RGI: Resistance Gene Identifier
RiPP: Ribosomally synthesized and post-translationally modified Peptide
RuBisCO: Ribulose-1,5-bisphosphate Carboxylase/Oxygenase
RTX-like: Repeats-in-Toxin–like
SoF1: *Xanthobacter* sp. SoF1 (strain designation)
TC-like: Toxin complex–like
TFBS: Transcriptor Factor Binding Sites
UniprotKB: UniProt Knowledgebase
VFDB: Virulence Factor Database

## Code availability

The source code, along with all other Python scripts and notebooks used to reproduce the data generated for this publication, are available on GitLab at https://git.rwth-aachen.de/hydrocow/pro-jects/comp-gen-so-f-1-vs-h-16.git

## Acknowledgements

The authors gratefully acknowledge the computing time provided to them at the NHR Center NHR4CES at RWTH Aachen University (project number rwth1662). This is funded by the Federal Ministry of Education and Research, and the state governments participating on the basis of the resolutions of the GWK for national high-performance computing at universities (https://www.nhr-verein.de/unsere-partner). The authors also acknowledge Solar Foods for providing the genome sequence of SoF1. Parts of the manuscript text were initially generated and later language-edited with the assistance of ChatGPT. All AI-assisted text was critically reviewed, revised, and verified by the authors, who take full responsibility for the accuracy, interpretation, and final content of the manuscript.

## Author information

### CRediT authorship contribution statement

K.K.: Conceptualization, Methodology, Software, Formal analysis, Investigation, Data curation, Writing – Original Draft, Writing - Review & Editing, Visualization. J-P.P.: Resources, Writing - Review & Editing, Project administration, Funding acquisition. L.M.B.: Writing - Review & Editing, Project administration, Funding acquisition. T.B.A: Conceptualization, Resources, Writing - Review & Editing, Supervision, Project administration, Funding acquisition

## Ethics declarations

### Ethics approval and consent to participate

Not applicable.

### Consent for publication

Not applicable.

